# Brain functional connectivity, but not neuroanatomy, captures the interrelationship between sex and gender in preadolescents

**DOI:** 10.1101/2024.10.31.621379

**Authors:** Athanasia Metoki, Roselyne J. Chauvin, Evan M. Gordon, Timothy O. Laumann, Benjamin P. Kay, Samuel R. Krimmel, Scott Marek, Anxu Wang, Andrew N. Van, Noah J. Baden, Vahdeta Suljic, Kristen M. Scheidter, Julia Monk, Forrest I. Whiting, Nadeshka J. Ramirez-Perez, Deanna M. Barch, Aristeidis Sotiras, Nico U.F. Dosenbach

**Author notes:** Correspondence to: Department of Neurology, Washington University School of Medicine in St. Louis, 660 S Euclid Ave, St. Louis, MO 63110, USA., E-mail address: athanasia (A. Metoki). Co-senior authors (Aristeidis Sotiras, Nico U.F. Dosenbach).

## Abstract

Understanding sex differences in the adolescent brain is crucial, as these differences are linked to neurological and psychiatric conditions that vary between males and females. Predicting sex from adolescent brain data may offer valuable insights into how these variations shape neurodevelopment. Recently, attention has shifted toward exploring socially-identified gender, distinct from sex assigned at birth, recognizing its additional explanatory power. This study evaluates whether resting-state functional connectivity (rsFC) or cortical thickness more effectively predicts sex and sex/gender alignment (the congruence between sex and gender) and investigates their interrelationship in preadolescents. Using data from the Adolescent Brain Cognitive Development (ABCD) Study, we employed machine learning to predict both sex (assigned at birth) and sex/gender alignment from rsFC and cortical thickness. rsFC predicted sex with significantly higher accuracy (86%) than cortical thickness (75%) and combining both did not improve the rsFC model’s accuracy. Brain regions most effective in predicting sex belonged to association (default mode, dorsal attention, and parietal memory) and visual (visual and medial visual) networks. The rsFC sex classifier trained on sex/gender aligned youth was significantly more effective in classifying unseen youth with sex/gender alignment than in classifying unseen youth with sex/gender unalignment. In females, the degree to which their brains’ rsFC matched a sex profile (female or male), was positively associated with the degree of sex/gender alignment. Lastly, neither rsFC nor cortical thickness predicted sex/gender alignment. These findings highlight rsFC’s predictive power in capturing the relationship between sex and gender and the complexity of the interplay between sex, gender, and the brain’s functional connectivity and neuroanatomy.

## 1. Introduction

The investigation of sex differences in the human brain has been a research interest for decades (Cosgrove et al., 2007; Giedd et al., 2012; Jäncke, 2018; Kaczkurkin et al., 2018; Lenroot & Giedd, 2010; Luders & Kurth, 2020; Luders & Toga, 2010; Sacher et al., 2013) spanning studies focusing on adults (Choleris et al., 2018; Jäncke, 2018; Kodiweera et al., 2016; Lotze et al., 2019; Ritchie et al., 2018), to those examining children and adolescents (Giedd et al., 2012; R. E. Gur & Gur, 2016; Herting & Sowell, 2017; Tunç et al., 2016; Vijayakumar et al., 2018), and even infants (Benavides et al., 2018; Fenske et al., 2023; Gilmore et al., 2007). Understanding brain sex differences is crucial, as many neurological and psychiatric conditions vary by sex in prevalence, onset, and symptomatology (Bao & Swaab, 2010; Baron-Cohen et al., 2011; Paus et al., 2008; Rutter et al., 2003), possibly due to differences in brain structure or function. Adolescence is a critical developmental period marked by increasing differences between males and females in physical traits, behavior, and mental health risks. Identifying sex-specific brain development patterns during this time could illuminate the roots of distinct mental health vulnerabilities, guiding tailored interventions that address the unique needs of each sex.

Diverse neuroimaging modalities have been used to study brain sex differences, including anatomical (e.g., volume, surface, and white matter integrity) and functional measures (e.g., resting-state functional connectivity; rsFC). Structural variations have been observed between males and females (for a comprehensive review see Kaczkurkin et al., 2018), in brain size (Ball et al., 2012; Bramen et al., 2011; De Bellis et al., 2001; Gennatas et al., 2017; Luders et al., 2005; Ritchie et al., 2018; Ruigrok et al., 2014; Sowell et al., 2007), cortical and subcortical volume (Allen et al., 2003; Goldstein et al., 2001; Gur et al., 1999; Luders et al., 2005), cortical surface area, hemispheric white matter connectivity (Ingalhalikar et al., 2014), and cerebral blood flow (Satterthwaite et al., 2014). Despite these structural distinctions, the existence and extent of functional brain organization sex differences remains unclear. While some studies have pinpointed specific networks where rsFC appears to vary between males and females (Biswal et al., 2010; Ritchie et al., 2018; L. Tian et al., 2011; Zuo et al., 2010), others have reported no discernible impact of sex in resting-state functional magnetic resonance imaging (fMRI) data (Weissman-Fogel et al., 2010). Moreover, evidence shows that sex differences in rsFC networks are affected by the menstrual cycle (Avila-Varela et al., 2024; Weis et al., 2019), indicating that rsFC variation may depend not only on sex but also on underlying factors like hormonal fluctuations.

Most studies on brain sex differences focus on sex as assigned at birth. However, an individual’s socially-identified gender may also influence brain development, potentially in part through societal and cultural perceptions of and reactions to their gender. This, in turn, can contribute to gender behavioral differences independent of sex (Bale & Epperson, 2016; Kim & Nafziger, 2000). To explore whether brain factors play a role in gender identity development, researchers have compared cisgender and transgender adult individuals, with mixed findings. Some studies suggest that the brains of transgender individuals resemble those of their sex assigned at birth (Emory et al., 1991; Luders et al., 2009; Mueller, Landré, et al., 2017; Yokota et al., 2005), others align them with their gender identity (Mueller, Landré, et al., 2017; Zhou et al., 1995), while additional research shows an intermediate(Flint et al., 2020; Kranz et al., 2014; Kurth et al., 2022; Mueller, De Cuypere, et al., 2017; Mueller, Landré, et al., 2017; Rametti, Carrillo, Gómez-Gil, Junque, Segovia, et al., 2011; Rametti, Carrillo, Gómez-Gil, Junque, Zubiarre-Elorza, et al., 2011) or unique phenotype (Mueller et al., 2021). Overall, evidence on brain differences related to sex and gender identity remains inconclusive.

Previous research on brain sex differences has often relied on traditional regression analyses that treat networks or regions-of-interest (ROI) independently. In contrast, multivariate pattern analysis can reveal more complex patterns without assuming independence, making advanced computational methods better suited for identifying brain sex differences. Machine learning approaches, such as classification, can assess how accurately sex can be predicted from neuroimaging data (Bzdok, 2017), yet few studies have applied these methods to examine sex differences in youth (Adeli et al., 2020; Brennan et al., 2021; Kurth et al., 2020; Sepehrband et al., 2018; Shanmugan et al., 2022). Existing research has typically focused on either brain structure (Adeli et al., 2020; Brennan et al., 2021; Kurth et al., 2020; Sepehrband et al., 2018) or function (Shanmugan et al., 2022), with no study to date comparing the predictive power of neuroanatomy versus functional connectivity in children or adolescents using a single dataset. Additionally, the potential benefits of combining both structural and functional data have not been explored in young populations, and most studies have used small samples spanning broad age ranges (Kurth et al., 2020; Sepehrband et al., 2018; Shanmugan et al., 2022) rather than focusing on larger, age-specific samples. Only one study has examined the neurobiological underpinnings of sex and gender in children (Dhamala et al., 2024) using linear ridge regression, finding that while sex and gender are uniquely represented in brain patterns, gender is less distinctly captured in functional connectivity than sex. However, this study did not explore a key cross-comparison, leaving unresolved questions about whether these unique representations share predictive power or are entirely independent.

In the present study, we moved beyond previous work by evaluating whether rsFC or cortical thickness is more effective in predicting sex and gender, while also exploring the extent of their interrelationship within the brain of preadolescents. We leveraged neuroimaging data from the Adolescent Brain Cognitive Development^®^ (ABCD; Garavan et al., 2018) Study (*n* = 3,196, 9-10 years old) at baseline, and self- and parent-reported gender data at the one-year follow-up time point. We use the following terminology: “sex” to describe an individual’s sex assigned at birth (i.e., male or female), usually based on physical anatomy and/or chromosomes at birth, and “gender” as the internal sense of oneself as boy, girl, or something else (Potter et al., 2021). We also use the term “sex/gender alignment” to refer to the congruence between an individual’s sex and their gender. Our approach involved unimodal (i.e., rsFC and cortical thickness separately) and multimodal (i.e., rsFC and cortical thickness combined) support vector machine (SVM) learning neuroimaging analyses to determine the predictability of sex and sex/gender alignment. Additionally, we tested whether a sex classifier trained on individuals with sex/gender alignment would perform comparably when predicting the sex of similarly aligned versus unaligned individuals. Lastly, we examined the relationship between the rsFC and cortical thickness sex classification scores (brain profile predictions) and sex/gender alignment scores to assess the consistency between the adolescents’ brain profiles and their sex/gender alignment.

## 2. Methods

### 2.1. ABCD study

The ABCD is a ten-year, longitudinal study of 11,875 youth enrolled at ages 9-10 from 21 sites in the United States. Participants (youth and parents) were recruited through schools, with minimal exclusion criteria (Garavan et al., 2018). The ABCD study obtained centralized institutional review board (IRB) approval from the University of California, San Diego. Each of the 21 sites also obtained local IRB approval. Ethical regulations were followed during data collection and analysis. Parents or caregivers provided written informed consent, and children gave written assent. The racial demographics of the participants roughly match the racial composition of the 2023 American Community Survey (U.S. Census Bureau, 2023). Although the ABCD study utilizes a longitudinal design, the analyses conducted for the current study were cross-sectional.

### 2.2. Participants

This project used rsFC and cortical thickness data from *n* = 10,259 available participants from the ABCD BIDS (Brain Imaging Data Structure) Community Collection (Feczko et al., 2021) (ABCD collection 3165; https://github.com/ABCD-STUDY/nda-abcd-collection-3165), baseline visit (9-10y). Demographic and behavioral data were obtained from the ABCD 4.0 release, which included data from the one-year follow-up visit, marking the beginning of gender measures’ collection. Following the ABCD consortium’s recommendations, we excluded participants scanned with Philips scanners due to incorrect preprocessing (source: https://github.com/ABCD-STUDY/fMRI-cleanup).

Head motion can systematically bias developmental studies (Power et al., 2012; Siegel et al., 2017), as well as those relating rsFC to behavior (Siegel et al., 2017). However, these systematic biases can be addressed through rigorous head motion correction (Power et al., 2014). The inclusion criteria for the current project required participants to have at least 600 frames (equivalent to 8 minutes) of low-motion (Fair et al., 2020) rsFC data (low-motion defined as having a filtered framewise displacement (FD) of less than 0.08 mm). See Casey et al. (2018) for broader ABCD inclusion criteria. Based on these criteria, and after excluding subjects with missing values in demographic, gender, or behavioral questions, the final ABCD sample consisted of rsFC, cortical thickness, demographic, gender, and behavioral data from a total of *n* = 3,196 youth (Table 1).

**Table 1.** Demographics (n = 3,196) and descriptive statistics. SD, standard deviation; PDS, pubertal development scale; FD framewise displacement.

| TOTAL (n = 3,196) |  |  |  |
| --- | --- | --- | --- |
|  | Males | Females |  |
|  | n = 1,573; 49.2% | n = 1,623; 50.8% |  |
| Mean Age in Months (SD) | 132.4 (7.8) | 132.3 (7.7) | t = 0.45, p = 0.65 |
| Range | 116-148 | 117-149 |  |
| Sex/Gender Alignment |  |  |  |
| Aligned | 1,172 (74.5%) | 873 (53.8%) | $\chi^2 = 148.79, p < 0.001$ |
| Unaligned | 401 (25.5%) | 750 (46.2%) |  |
| PDS |  |  |  |
| Prepuberty | 476 (30.3%) | 226 (13.9%) | $\chi^2 = 124.37, p < 0.001$ |
| Early puberty | 727 (46.2%) | 337 (20.7%) | $\chi^2 = 233.03, p < 0.001$ |
| Mid puberty | 350 (22.2%) | 873 (53.8%) | $\chi^2 = 336.35, p < 0.001$ |
| Late puberty | 20 (1.3%) | 188 (11.6%) | $\chi^2 = 138.56, p < 0.001$ |
| Residual In-Scanner | 0.13 (0.08) | 0.12 (0.07) | $t = 3.79, p < 0.001$ |
| Motion (Mean FD), mm | 0.03-1.07 | 0.03-0.70 |  |

### 2.3. MRI acquisition and image processing methods

#### 2.3.1. MRI acquisition

Imaging for each ABCD youth was performed across 21 sites within the United States, and was harmonized across Siemens Prisma, Philips, and GE 3T scanners. Details on image acquisition can be found elsewhere (Casey et al., 2018). Twenty minutes (4 × 5 min runs) of eyes-open (passive crosshair viewing) resting-state fMRI data were collected to ensure at least 5 minutes of low-motion data. All resting-state fMRI scans were acquired using a gradient-echo EPI sequence (TR = 800ms, TE = 30ms, flip angle = 90°, voxel size = 2.4mm^3^, 60 slices). Head motion was monitored online using the Framewise Integrated Real-time MRI Monitor (FIRMM) software at Siemens sites (Dosenbach et al., 2017).

#### 2.3.2. Image processing overview

The image processing stream has been detailed elsewhere (Feczko et al., 2021). Briefly, the ABCD pipeline comprises six stages: (1) PreFreeSurfer, which normalizes anatomical data; (2) FreeSurfer, which constructs cortical surfaces from the normalized anatomical data; (3) PostFreeSurfer, which converts outputs from FreeSurfer to CIFTIs and transforms the volumes to a standard volume space using ANTs nonlinear registration; (4) “Vol” stage, which performs the atlas transformation, mean field distortion correction, and resampling to 2 mm isotropic voxels in a single step using FSL’s applywarp tool; (5) “Surf” stage, which projects the volumetric functional data onto the surface; and (6) “DCANBOLDproc”, which performs functional connectivity processing.

DCANBOLDproc includes a respiratory filter to improve FD estimates calculated in the “vol” stage. Temporal masks were created to flag motion-contaminated frames using the improved FD estimates (Power et al., 2012). Frames with FD > 0.30 mm were flagged as motion contaminated. After computing the temporal masks for high motion frame censoring, the data were processed with the following steps: (i) demeaning and detrending, (ii) interpolation across censored frames using least squares spectral estimation of the values at censored frames (Power et al., 2014) so that continuous data can be (iii) denoised via a general linear model (GLM) including whole brain, ventricular, and white matter signals, as well as their derivatives.

Denoised data were then passed through (iv) a band-pass filter (0.008 Hz < f < 0.10 Hz) without re-introducing nuisance signals (Hallquist et al., 2013) or contaminating frames near high motion frames (Carp, 2013).

#### 2.3.3. Functional connectivity and cortical thickness metrics

An ROI method was employed, utilizing a total of 394 parcels, which included the 333 cortical parcels initially defined by Gordon et al. (2016) and an additional 61 subcortical parcels described by Seitzman et al. (2020). For each participant, the rsFC timecourse was extracted from these 394 ROIs after discarding frames with a filtered FD > 0.08 mm. The 333 cortical ROIs can be organized into separable brain networks (e.g., default-mode, fronto-parietal, salience, etc.). A correlation matrix was created through the computation of the correlation between the timecourse of each ROI and that of every other ROI, yielding a 394×394 correlation matrix for each subject. The correlations were normalized using Fisher’s r-to-z transform (Fisher, 1915). The correlation matrices were then combined across all subjects to form a 394×394×3,196 matrix, which was employed for subsequent analyses. rsFC analyses were run at the edge level (ROI-ROI pair; *n* = 77,421). Furthermore, cortical thickness was extracted from the 333 cortical ROIs for each participant. Additional cortical thickness analyses at the vertex-level (59,412 cortical vertices for each participant) were also conducted in an effort to capture regional effects that may not align with ROI boundaries (Supplementary Results).

### 2.4. Sex and gender measures

We used the ABCD’s parent-reported data on youths’ sex, which reflect their sex assigned at birth. The ABCD study assesses gender with the Youth Self-Report and Parent-Report Gender Questionnaires (Supplementary Table 1). All participants included in analyses completed the Youth Self-Report Gender Questionnaire which comprised four questions assessing felt-gender, contentedness with assigned sex at birth, and gender expression. Their parents/caregivers completed an adaptation of the Gender Identity Questionnaire (Elizabeth & Green, 1984; Johnson et al., 2004) which included 12 questions that measure sex-typed behavior during play and gender dysphoria (Potter et al., 2022). All items use a five-point scale with higher scores reflecting more congruence with sex assigned at birth. The current study computed and used the mean scores of both constructs separately, capturing gender dimensionally (Potter et al., 2022). Sex/gender alignment status was either “aligned”, i.e., sex/gender alignment defined as scoring the highest possible achievable score in the Youth Self-Report Gender Questionnaire, or “unaligned”, i.e., sex/gender unalignment defined as scoring anything other than the highest possible score. Participants who provided a “Decline to answer” response to any question were not included in the analyses (see Supplementary Figure 1 for response frequencies on the Youth Self-Report and Parent-Report Gender Questionnaires).

### 2.5. Support vector machine learning

#### 2.5.1. Support vector machine classification

Support vector machine (SVM) learning was used to examine the predictability of sex in adolescents based on neuroimaging measures. Three binary linear SVM classifiers were built: one with rsFC, one with cortical thickness, and one with rsFC and cortical thickness combined as predictors/features. The target labels for all models were the sex categories of male and female. The classifiers were trained exclusively using youth with sex/gender alignment (highest possible attainable score in the Youth Self-Report Gender Questionnaire). The aligned group data was split to retain 20% of subjects held out entirely for testing and 80% for training. Depending on the analysis, independent testing sets were composed either entirely of youth with sex/gender alignment (aligned group) or entirely of youth with sex/gender unalignment (unaligned group) (Fig. 1). To account for statistically significant sex differences in pubertal development stage and residual in-scanner motion (mean FD) (see Other Statistical Analyses, Results, & Table 1), these variables were regressed out separately from each feature in the training, cross-validated, and independent test datasets to prevent data leakage. To ensure robust model evaluation and prevent overfitting, the classification was evaluated by using a nested five-fold cross-validation (5F-CV) (Poldrack et al., 2020) in the training set, with the inner 5F-CV determining the optimal regularization coefficient C via grid search for the SVM binary classifier and the outer 5F-CV estimating the generalizability of the model (Fig. 1). Each feature was linearly scaled between zero and one across the outer 5F-CV training sets; these scaling parameters were then applied to scale the outer 5F-CV and independent testing sets (Cui & Gong, 2018; Erus et al., 2015). For additional SVM parameters see Supplementary Methods.

**Figure 1.**
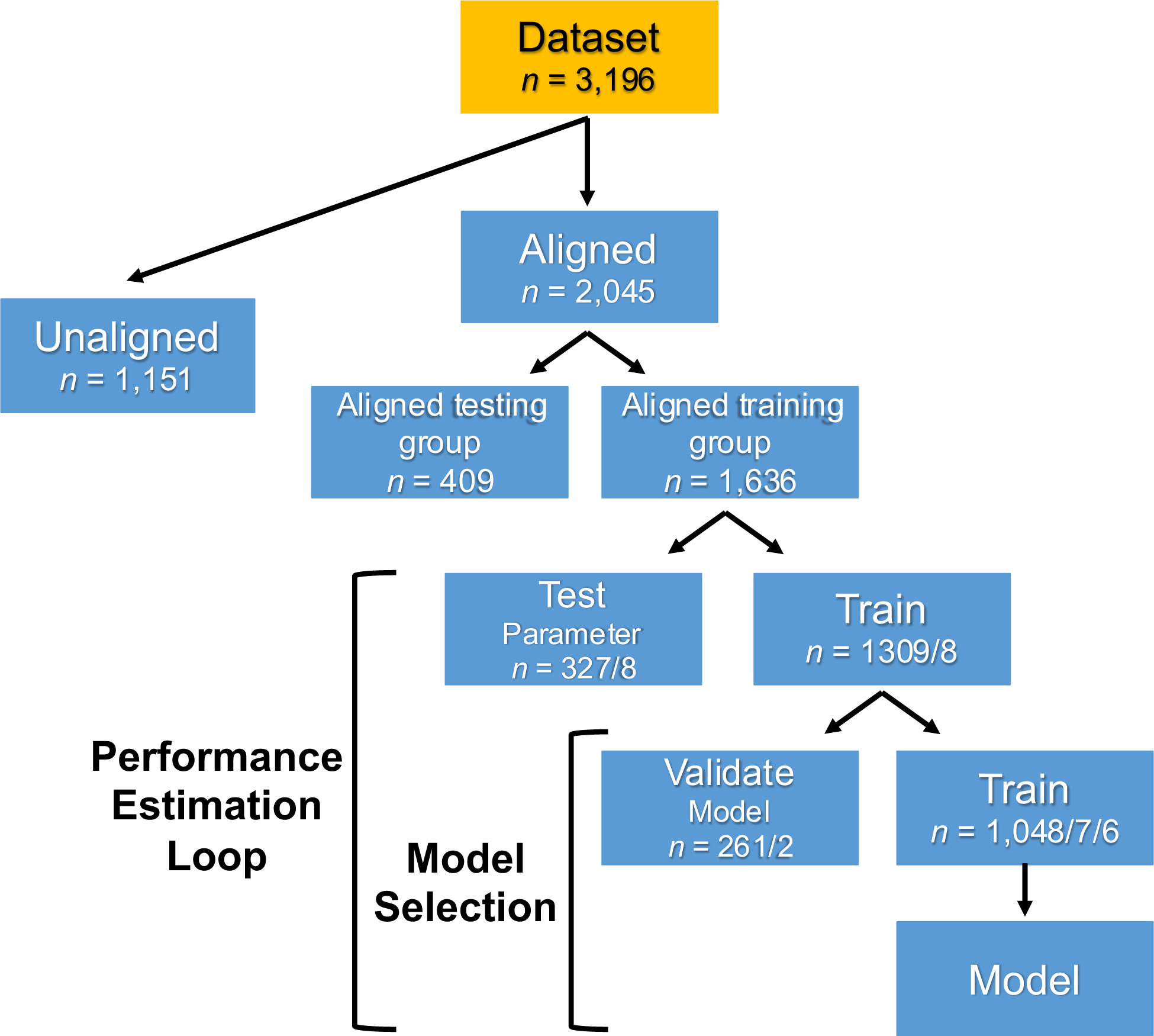
Flowchart of the support vector machine (SVM) model construction. The dataset was split in two groups: Aligned (participants with sex/gender alignment) and unaligned (participants with sex/gender unalignment). The aligned group was split in an aligned training group (80%) and an aligned hold-out testing group (20%). A nested five-fold cross-validation (5F-CV) was employed, with the inner 5F-CV determining the optimal parameter C and the outer 5F-CV estimating the generalizability of the model. The final, optimal model was subsequently applied in the held-out aligned testing group and the unaligned group and model performance was evaluated. The forward slash in the groupings denotes the different possible group splits (e.g. The aligned training group was split in four test/train group pairs with 327/1903 subjects and one test/train group pair with 328/1308 subjects).

Assigning weights to observations (single data points or instances within a dataset) is a valuable approach for addressing the issue of treating all observations equally. The weights indicate the importance or influence of each observation during the SVM model training. Due to an imbalanced distribution of males and females in the aligned group (see Results & Table 1), which would lead to an imbalance in the SVMs’ aligned training groups, inverse proportion instance weights were calculated and assigned to help balance the contribution of each class to the models.

#### 2.5.2. Support vector regression

Linear support vector regression (SVR) was used to evaluate whether sex/gender alignment (a discrete variable derived from the Youth Self-Report and Parent-Report Gender Questionnaires) can be predicted by brain neuroanatomy or functional connectivity. Using the same parameters/choices as in the SVM classification models (Methods & Supplementary Methods), we ran a total of twelve models, utilizing different combinations of features and questionnaire types. For the rsFC analyses, two models were run: one exclusive to females and one exclusive to males, and we employed a data split of 20% for hold-out testing and 80% for training, in both the female and male datasets. Each of these models utilized the Youth Self-Report Gender Questionnaire as the target variable in one iteration and the Parent-Report Gender Questionnaire in another. Similarly, for the cortical thickness and the combined rsFC/cortical thickness analyses, another set of models was executed, following the same structure as the rsFC analyses. The same grid search optimization of the regularization coefficient C using 5F-CV was performed as for the binary SVM classifiers (Supplementary Methods). Each feature was linearly scaled between zero and one across the outer 5F-CV training sets and these scaling parameters were then applied to scale the outer 5F-CV and the independent testing sets (Cui & Gong, 2018; Erus et al., 2015). The imbalance in the distribution of test scores in the Youth Self-Report and the Parent-Report Gender Questionnaires (Supplementary Figure 1) led to assigning inverse proportion instance weights in the SVR models.

As a robustness check measure for the rsFC SVR analyses, we opted to run linear ridge regressions as performed by Dhamala et al. (2024) using the code they provided online (https://zenodo.org/doi/10.5281/zenodo.10779163). The only adjustment in the script was to use the updated “StandardScaler” to normalize the data, as the “normalize” parameter has been removed from the Scikit-learn (Pedregosa et al., 2011) “Ridge” regression command starting from version 1.0, recommending users handle normalization separately instead.

#### 2.5.3. Assessment and significance of prediction performance

The performance evaluation of the SVM binary classification models encompassed a comprehensive set of metrics (Poldrack et al., 2020), including accuracy, sensitivity, specificity, the area under the receiver operating characteristic curve (AUC), and the Matthews correlation coefficient (MCC). As opposed to accuracy, which can produce overoptimistic inflated results (Akosa, 2017; Flach & Kull, 2015; Gu et al., 2009; Hand & Christen, 2018; Sokolova et al., 2006; Xi & Beer, 2018), the MCC treats both classes symmetrically, penalizing models that perform well on one class but poorly on another, thus providing a balanced evaluation (Chicco, Tötsch, et al., 2021; Chicco, Warrens, et al., 2021; Chicco & Jurman, 2020, 2022, 2023). It ranges from −1 to +1, with +1 indicating a perfect prediction, 0 a prediction no better than random chance, and −1 total disagreement between prediction and observation. This comprehensive suite of metrics ensured a thorough and nuanced evaluation of the binary classification models, capturing various aspects of their performance and facilitating informed decision-making.

The McNemar’s test (McNemar, 1947; Pembury Smith & Ruxton, 2020) was used to evaluate the differences in classification accuracies between the rsFC, cortical thickness, and combined rsFC/cortical thickness SVM binary classification models. A two-proportions *z*-test was conducted to compare the proportions of correct predictions between the aligned and the unaligned SVM models.

Permutation testing was used to evaluate whether the prediction performance of the SVM binary classification models was significantly better than expected by chance (Mourão-Miranda et al., 2005). The predictive framework was repeated 10,000 times. The coefficient of determination (*R^2^*) (Poldrack et al., 2020) was used as a method of evaluation of the SVR models’ performance.

#### 2.5.4. Interpreting model feature weights

The transformation proposed by Haufe et al. (2009) was utilized to transform feature weights derived from the linear SVM binary classification models with the aim of enhancing their interpretability and reliability (Chen et al., 2022; Y. Tian & Zalesky, 2021). To enhance interpretability for visualization purposes, the parcel-wise feature weights were consolidated at the network level by calculating their root mean square value and averaging across previously defined canonical functional networks (Gordon et al., 2016, 2023). Subcortical and cerebellar parcels were grouped into a single non-cortical network.

### 2.6. Other Statistical Analyses

To ensure that there were no confounders when running the SVM classification, we conducted a series of statistical tests to examine potential differences between males and females, allowing for a more accurate and unbiased analysis. Independent *t*-tests were utilized to compare age and residual in-scanner motion (mean FD) between the two groups. Additionally, chi-squared tests were applied to assess the distribution of males and females across various categorical variables, including sex/gender alignment, pubertal development stage, and family income.

The National Academies of Science, Engineering, and Medicine (NASEM) issued a report providing recommendations on using race, ethnicity, ancestry, and other descriptors of population stratification (National Academies of Sciences et al., 2023). The NASEM report asserts that race does not have any biological basis and emphasizes that researchers should not “control for race” in brain-based association or prediction studies with non-brain variables. The current study follows these recommendations, and therefore does not evaluate race and ethnicity in brain association and prediction models (Bird & Carlson, 2024; Schraiber & Edge, 2024).

Spearman’s correlations were used to examine the relationship between the rsFC and cortical thickness sex classification scores (brain profile predictions) and sex/gender alignment scores in order to evaluate the consistency between the adolescents’ brain profiles and sex/gender alignment. We opted for a non-parametric approach due to the non-normal distribution of the sex/gender alignment scores (Supplementary Figure 1).

## 3. Results

### 3.1. Youths’ demographic characteristics

In this sample, *n* = 3,196, males (*n* = 1,573 [49.2%]) and females (*n* = 1,623 [50.8%]) did not differ statistically in age (Table 1) or family income (Supplementary Table 2). Statistically significant sex differences were found in pubertal development scale scores for each puberty stage and residual in-scanner motion (mean FD of retained fMRI frames) (Table 1). Thus, pubertal development stage and mean FD, were used as covariates in all SVM analyses.

### 3.2. Functional connectivity and neuroanatomical classification of sex

This study’s first goal was to understand the way in which high dimensional patterns of brain functional connectivity and neuroanatomy reflect sex. The rsFC sex classifier trained on the aligned participants was able to classify unseen aligned participants from the independent testing set as male or female with 86% accuracy (*p* < 0.001; Fig. 2A, 2B). Sensitivity, specificity, AUC, and MCC of the model were 0.84, 0.89, 0.93, and 0.71 respectively. Variations in the functional organization of the visual (visual and medial visual), default mode, dorsal attention, and parietal memory networks, in that order, contributed the most to the model and were therefore relatively more important in predicting participant sex (Fig. 2C, 2D).

**Figure 2.**
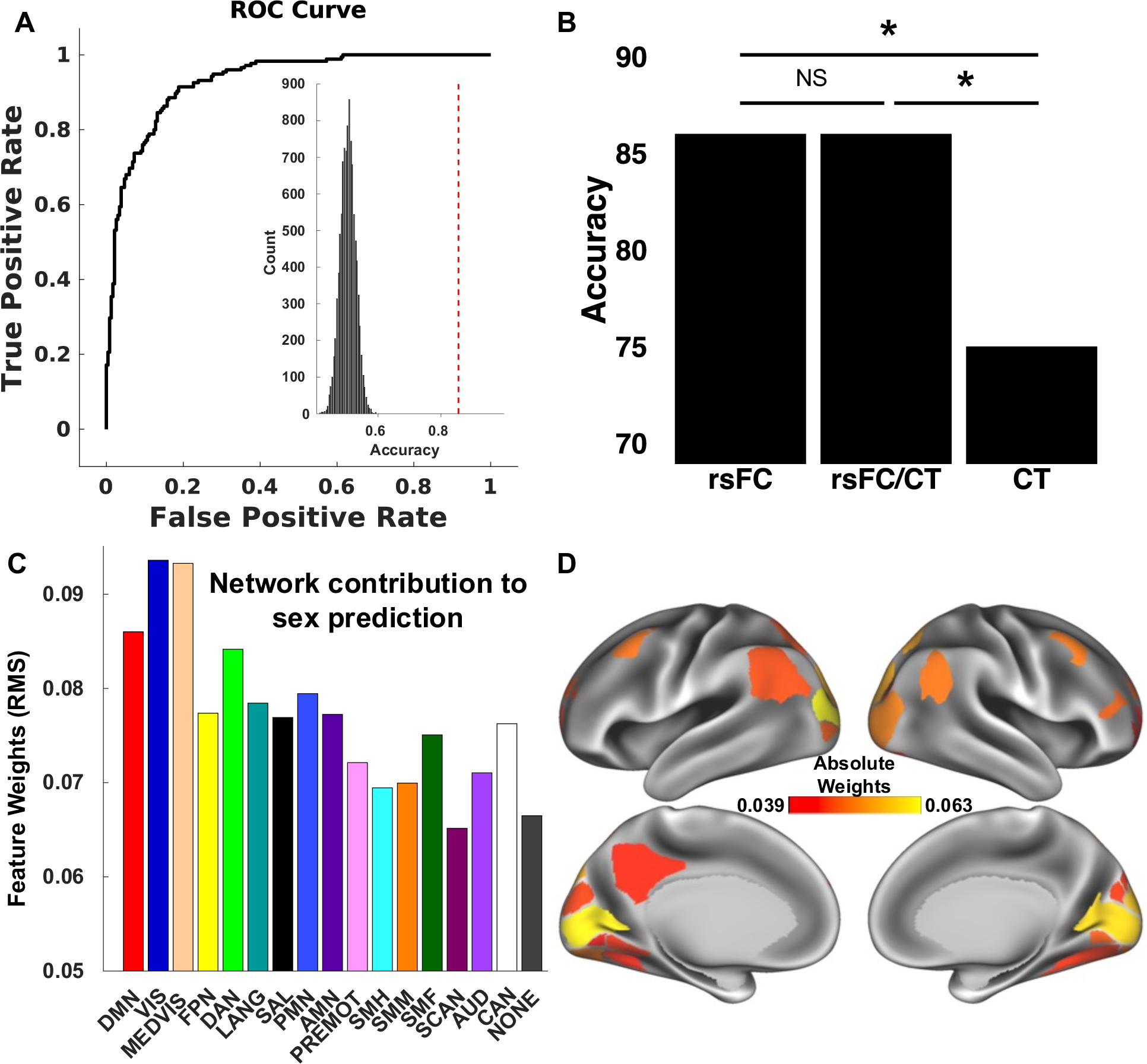
Brain pattern analysis using support vector machine (SVM) learning predicts participant sex based on functional connectivity. (A) SVMs with nested five-fold cross-validation (5F-CV) were used to construct univariate and multivariate models that classified participants as male or female (sex assigned at birth). These models used resting-state functional connectivity (rsFC), cortical thickness, and rsFC/cortical thickness combined as predictive features. The receiver operating characteristic (ROC) curve of the rsFC resulting model is depicted. The model classified participants as male or female with 86% accuracy. Inset histogram shows distribution of permuted accuracies. The accuracy from real (nonpermuted) data is represented by the dashed red line. (B) McNemar’s tests comparing the rsFC, cortical thickness, and combined rsFC/cortical thickness sex classifiers revealed a statistically significant difference between the rsFC and cortical thickness performance (*χ^2^*= 10.96, *p <* 0.001) and between the combined rsFC/cortical thickness and cortical thickness performance (*χ^2^* = 10.96, *p <* 0.001), but a non-significant difference between the rsFC and combined rsFC/cortical thickness performance (*χ^2^* = 0.01, *p* = 93). (C) To understand which networks contributed the most to the prediction, the parcel-wise feature weights’ root mean square was calculated and then averaged for each network. The most important features in the rsFC model were found in the visual (visual and medial visual), default mode, dorsal attention, and parietal memory networks. (D) The top 10% of cortical parcels in terms of feature importance in the rsFC SVM model. The parcel-wise feature weights’ root mean square was calculated for each parcel. rsFC, resting-state functional connectivity; CT, cortical thickness; NS, not significant; DMN, default mode network; VIS, visual network; MEDVIS, medial visual network; FPN, frontoparietal network; DAN, dorsal attention network; LANG, language network; SAL, salience network; PMN, parietal memory network; AMN, action-mode network; PREMOT, premotor network, SMH, somatomotor hand network; SMM, somatomotor mouth network; SMF, somatomotor foot network; SCAN, somato-cognitive action network; AUD, auditory network; CAN, contextual association network; NONE, subcortical and cerebellar structures.

The cortical thickness sex classifier was trained on the same aligned participants as the rsFC classifier. It correctly separated aligned males from females with an accuracy of 75% (*p* < 0.001; Fig. 2B). Sensitivity, specificity, AUC, and MCC of the model were 0.75, 0.75, 0.82, and 0.50 respectively. Variation in the anatomical organization of cortical parcels belonging to the medial visual, frontoparietal, visual, action mode, and dorsal attention networks, in that order, contributed the most to the model and were therefore relatively more important in predicting participant sex.

The combined rsFC/cortical thickness sex classifier was similarly trained on the same set of aligned participants. It correctly separated aligned males from females with an accuracy of 86% (*p* < 0.001; Fig. 2B). Sensitivity, specificity, AUC, and MCC of the model were 0.83, 0.89, 0.93, and 0.71 respectively. Variations in the organization of the visual (medial visual and visual), default mode, dorsal attention, and frontoparietal networks, in that order, contributed the most to the model and were therefore relatively more important in predicting participant sex.

The rsFC sex classifier exhibited significantly superior performance in predicting aligned individuals compared to the cortical thickness classifier (McNemar’s test: *χ^2^* = 10.96, *p <* 0.001; Fig. 1B). The combined rsFC/cortical thickness model also performed significantly better than the cortical thickness model (*χ^2^* = 10.96, *p <* 0.001), but there was no significant difference between the rsFC and the combined rsFC/cortical thickness classifier performances (*χ^2^* = 0.01, *p* = 93).

### 3.3. Sex classifiers’ efficacy in predicting aligned and unaligned individuals

Although the rsFC classifier demonstrated superior predictive performance for sex within the aligned test group, surpassing the cortical thickness classifier, we examined both and the combined rsFC/cortical thickness classifier to assess efficacy in predicting the sex of individuals with unaligned sex/gender. The rsFC sex classifier trained on the aligned participants was able to classify unseen unaligned participants as male or female with 79% accuracy (*p <* 0.001). Sensitivity, specificity, AUC, and MCC of the model were 0.85, 0.75, 0.82, and 0.58 respectively. The rsFC SVM model predicting the independent aligned group achieved statistically significantly higher prediction accuracy (86%) than predicting the unaligned group (79%; *z* = 3.08, *p* < 0.01).

The cortical thickness sex classifier trained on the aligned participants was able to classify unseen unaligned participants as male or female with 71% accuracy (*p <* 0.001; Sensitivity, specificity, AUC, MCC = 0.81, 0.65, 0.82, and 0.44). However, the cortical thickness SVM model did not achieve a statistically significantly higher prediction accuracy for the aligned independent testing set (75%) compared to the unaligned group (71%; *z* = 1.64, *p* = 0.10).

The combined rsFC/cortical thickness classifier sex classifier trained on the aligned participants was able to classify unseen unaligned participants as male or female with 78% accuracy (*p <* 0.001; Sensitivity, specificity, AUC, MCC = 0.86, 0.74, 0.89, and 0.57). Similar to the rsFC sex classifier, the rsFC/cortical thickness SVM model achieved a statistically significantly higher prediction accuracy for the aligned independent testing set (86%) compared to the unaligned group (78%; *z* = 3.21, *p* < 0.01).

Just as for aligned individuals, the rsFC sex classifier exhibited significantly superior performance in predicting unaligned individuals compared to the cortical thickness classifier (McNemar’s test: *χ^2^*= 13.28, *p <* 0.001). Similarly, the combined rsFC/cortical thickness sex classifier performed significantly better at predicting unaligned individuals compared to the cortical thickness classifier (McNemar’s test: *χ^2^*= 12.02, *p <* 0.001) but there was no significant difference between the rsFC and combined rsFC/cortical thickness classifier performances (*χ^2^* = 0.02, *p* = 89) in unaligned individuals.

Since combining rsFC and cortical thickness did not improve the rsFC classification model, the separate rsFC and cortical thickness models were used for the remainder of the classification-related analyses.

### 3.4. Sex classification and sex/gender alignment correlation

The above findings indicate robust differences in brain functional connectivity (rsFC) and neuroanatomy (cortical thickness) between males and females. The relationship between the rsFC and cortical thickness sex classification scores (brain profile predictions) and sex/gender alignment scores was then investigated to evaluate the consistency between the adolescents’ brain profiles and sex/gender alignment. The classification scores derived from the rsFC and cortical thickness sex classifiers serve as quantitative indicators of the degree to which a brain matches either female or male patterns. Higher scores suggest a stronger correspondence to the neural connectivity patterns associated with the sex commonly labeled as either female or male. Α significant positive correlation was revealed in females between the rsFC classification testing set scores and the sex/gender alignment scores obtained from the Youth Self-Report Gender Questionnaire (*ρ* = 0.17, *p <* 0.001, Fig. 3A). This suggests that the females with functional connectivity patterns more similar to the typical female pattern exhibit greater sex/gender alignment as reflected in higher scores on the Youth Self-Report Gender Questionnaire. In males, the inverse association was observed, with individuals with functional connectivity patterns more similar to the typical male pattern exhibiting lower sex/gender alignment *(ρ* = −0.16, *p* < 0.001; Fig. 3B).

**Figure 3.**
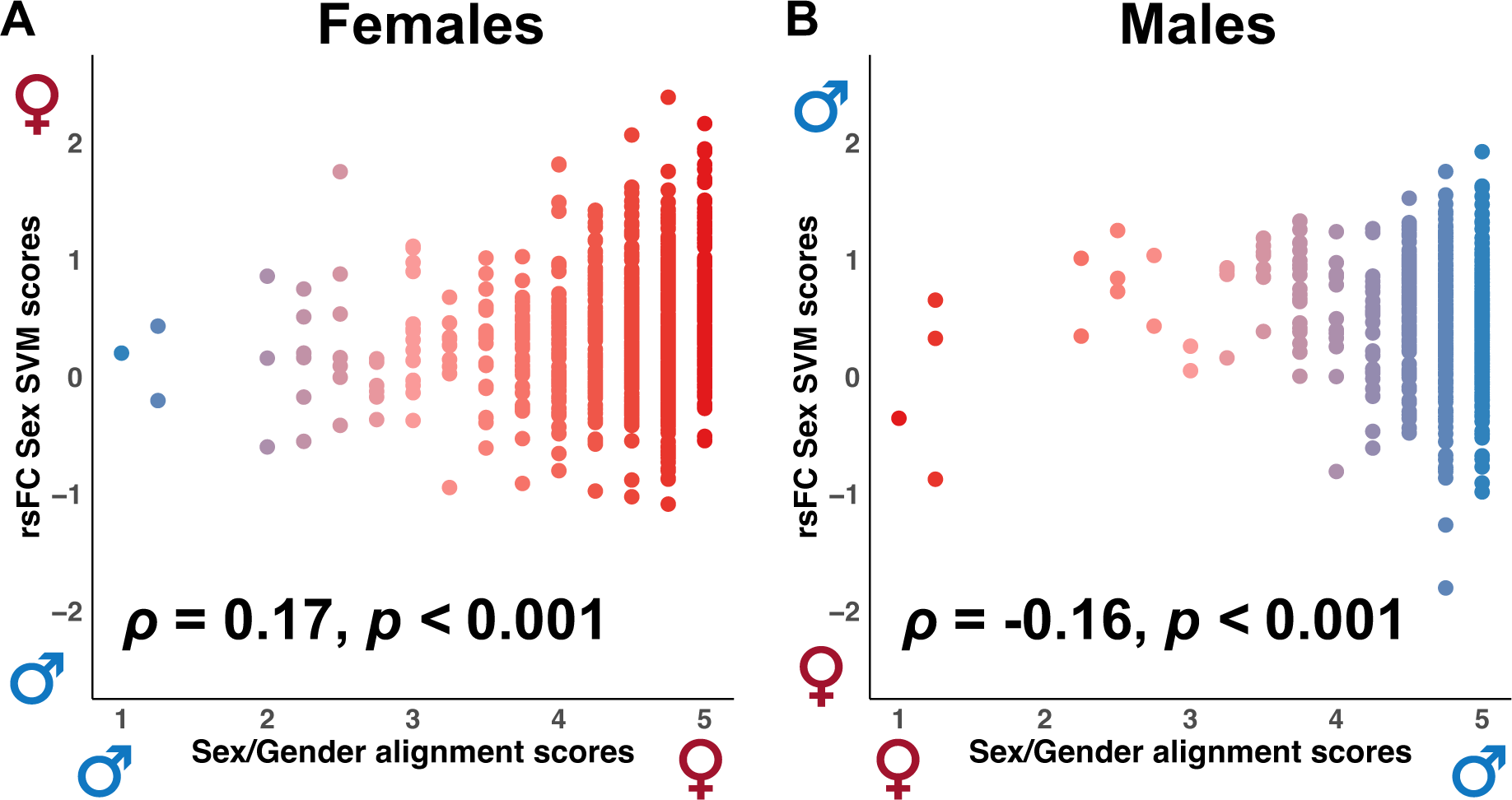
Correlations between resting-state functional connectivity (rsFC) support vector machine (SVM) classification scores and sex/gender alignment scores. (A) A significant positive correlation was observed in females between the rsFC sex classification scores and the sex/gender alignment scores. (B) A significant negative correlation between the rsFC classification and sex/gender alignment scores was observed in males. Higher sex/gender alignment scores (x-axis) indicate greater sex/gender alignment. Higher SVM scores (y-axis) indicate a stronger correspondence to the neural connectivity patterns associated with the sex labeled as either female or male. Results in bold indicate statistical significance.

The correlation between cortical thickness classification and sex/gender alignment scores was not significant for females (*ρ* = 0.04, *p* = 0.27) but was significant for males (*ρ* = −0.13, *p* < 0.001) exhibiting the same inverse relationship as with the rsFC brain patterns.

### 3.5. Functional connectivity and neuroanatomical prediction of sex/gender alignment

Next, we evaluated whether an adolescent’s degree of sex/gender alignment can be predicted by brain neuroanatomy or functional connectivity using linear SVR models. Neither the functional connectivity (rsFC), neuroanatomy (cortical thickness), nor the combined rsFC/cortical thickness sex/gender alignment SVR models successfully predicted the sex/gender alignment Youth Self-Report or Parent-Report Gender Questionnaire scores. Correlations between the original sex/gender alignment scores and predicted sex/gender alignment scores were not significant (all *p*-values > 0.05). Importantly, all coefficients of determination were negative (*R^2^* < 0) suggesting that the models performed worse than a null model predicting the mean of the dependent variable (sex/gender alignment scores) for all observations.

As a robustness check measure, linear ridge regressions, as performed by Dhamala et al. (2024) to explore functional brain correlates of sex and gender, were also tested. The findings were the same as in our rsFC SVR model. Correlations between the original Youth Self-Report and Parent-Report Gender Questionnaire gender scores and predicted gender scores were not significant (all *p*-values > 0.05) and all coefficients of determination were negative (*R^2^* < 0).

## 4. Discussion

In this study, we leveraged machine learning and a large sample of youths to determine whether variation in functional connectivity and neuroanatomy can predict sex (assigned at birth) and/or sex/gender alignment, and whether this variability is reflected in the degree of sex/gender alignment. Using complex univariate and multivariate patterns of rsFC and cortical thickness, we were able to predict an unseen participant’s sex with a high degree of accuracy. We first demonstrated that rsFC is a better predictor of sex compared to cortical thickness. Combining rsFC and cortical thickness did not improve the rsFC model’s performance. We then showed that sex differences in brain functional connectivity and neuroanatomy are greatest in association networks. Moreover, SVM analyses revealed that the rsFC of sex/gender aligned youth is more effective at predicting an unseen aligned participant’s sex compared to predicting the sex of an unaligned participant, while the cortical thickness SVM classifier failed to distinguish between aligned and unaligned participants. Furthermore, we found a positive relationship between the degree to which an individual’s rsFC profile reflects male or female traits and sex/gender alignment only in females. In males, we observed a pattern of negatively correlated relationships both in rsFC and cortical thickness. Lastly, we found that neither functional connectivity nor neuroanatomy predict sex/gender alignment.

### 4.1. rsFC outperforms cortical thickness as a predictor of sex

One of our principal findings is that rsFC demonstrates superior predictive power compared to cortical thickness in accurately classifying sex, although both modalities showed a good degree of prediction. Prior studies have demonstrated similar accuracies using network-level resting-state fMRI data to discriminate between male and female youth (Shanmugan et al., 2022), achieving slightly higher accuracies when combined with anatomical data (Adeli et al., 2020; Brennan et al., 2021; Kurth et al., 2020; Sepehrband et al., 2018). In our analysis, however, combining rsFC and cortical thickness was better than cortical thickness alone, but did not show a significant improvement in prediction accuracy compared to the rsFC-only model. The superior predictive power of rsFC suggests that, even in a young cohort, neural activity patterns during rest capture subtle, sex-specific functional dynamics that are not evident in structural measures like cortical thickness. This highlights rsFC’s potential as a more sensitive biomarker in neurodevelopmental studies. Additionally, the fact that rsFC outperforms cortical thickness as a predictor could imply that the information provided by cortical thickness overlaps with what is already captured by rsFC. As a result, cortical thickness may not offer additional or unique information to enhance prediction accuracy.

### 4.2. Sex differences in brain functional connectivity and neuroanatomy are greatest in association networks

Our findings reveal significant sex-related variability in brain functional connectivity and neuroanatomy, particularly within association networks, an observation consistent with prior research (Bijsterbosch et al., 2018; Cui et al., 2020; Gordon et al., 2017; Kong et al., 2019; Li et al., 2017; Shanmugan et al., 2022). The most pronounced differences in rsFC between sexes were found in visual, default mode, dorsal attention, and parietal memory networks (Filippi et al., 2013; Mathew et al., 2020; Murray et al., 2018; Shaqiri et al., 2018; Vanston & Strother, 2017; Weis et al., 2019). Studies suggest that men and women may process visual information differently (Mathew et al., 2020; Murray et al., 2018; Shaqiri et al., 2018; Vanston & Strother, 2017), though neuroimaging research in adults presents mixed results – some identify sex differences in sensory networks like the visual network (Biswal et al., 2010; Filippi et al., 2013), while others do not (Allen et al., 2011; Tomasi & Volkow, 2012). The functional organization of association networks, which are linked to emotional, social, and executive functions (Cui et al., 2020; Kong et al., 2019), also demonstrates variability tied to sex differences (Alarcón et al., 2020; Andreano & Cahill, 2009; Cahill, 2010). For instance, variations in default mode network connectivity have been associated with sex hormones such as estrogen and progesterone (Engman et al., 2016; Hidalgo-Lopez et al., 2020; Petersen et al., 2014; Weis et al., 2019). Our findings further support the role of the dorsal attention and parietal memory networks in predicting sex differences, which also corresponds with prior research (Dumais et al., 2018; Filippi et al., 2013; Hill et al., 2014; Hjelmervik et al., 2014; Stumme et al., 2020; Zhang et al., 2018). These sex differences in rsFC could relate to variations in cognitive processes and behavioral outcomes observed between males and females, influencing domains such as decision-making, emotion regulation, and attention allocation (Stoica et al., 2021). Our results partially align with the work of Dhamala et al. (2024), who also utilized the ABCD dataset to explore functional brain correlates of sex and gender, finding that sex is strongly associated with connectivity patterns within and between the somatomotor, visual, control, and limbic networks (Yeo et al., 2011). However, it is also possible that societal mores that influence engagement with certain cognitive activities also shape brain functional connectivity, leading to patterns that differ between sexes that are not biologically determined.

### 4.3. rsFC of sex/gender aligned youth is less accurate in predicting unaligned youths’ sex

We found that that observed differences in rsFC between males and females may not entirely be driven by biology (sex) but also from the influence of social constructs (gender) (Eliot, 2011; Eliot et al., 2021). Using a machine learning neuroimaging approach, we showed that an SVM classifier trained to learn sex from rsFC data using only youth with sex/gender alignment, was significantly more accurate in predicting sex in unseen participants with sex/gender alignment than in those with sex/gender unalignment. These findings suggest that there may be subtle yet detectable differences in rsFC patterns between individuals with aligned and unaligned sex and gender. Notably, a cortical thickness SVM classifier trained to learn sex using only youth with sex/gender alignment failed to differentiate effectively, as it was equally accurate in predicting sex in unseen participants with sex/gender alignment and those with unalignment. This might imply that gender has greater influence on functional connectivity brain patterns than on brain neuroanatomy (i.e., cortical thickness). These results could be considered distinct from research suggesting that the structural brain profile of hormonally untreated transgender individuals resembles that of their gender identity or is intermediate between sexes (Flint et al., 2020; Kranz et al., 2014; Kurth et al., 2022; Mueller, De Cuypere, et al., 2017; Mueller, Landré, et al., 2017; Rametti, Carrillo, Gómez-Gil, Junque, Segovia, et al., 2011; Rametti, Carrillo, Gómez-Gil, Junque, Zubiarre-Elorza, et al., 2011). However, those studies reported on white matter microstructure and gray matter volume, including subcortical structures, and not cortical thickness.

The reduced accuracy of the rsFC classifier when applied to gender-diverse youth highlights that the neurobiological patterns linked to sex in youth with sex/gender alignment do not generalize well to those with sex/gender unalignment. This underscores the need to consider gender diversity in neuroscientific research and cautions against relying on brain-based classifiers trained exclusively on sex/gender aligned individuals.

### 4.4. Female youth with more sex-specific brain patterns exhibit greater sex/gender alignment

We found that females with higher rsFC sex SVM scores (greater brain “femaleness” on the “maleness-femaleness” continuum) also had higher sex/gender alignment scores on the Youth Self-Report Gender Questionnaire (Fig. 3A). In males, however, there was a surprising negative correlation: males with higher rsFC sex SVM scores (greater brain “maleness”) showed lower sex/gender alignment (Fig. 3B). One possible explanation for this discrepancy may be linked to lower social tolerance for gender nonconformity in young males compared to females (Potter et al., 2021; Potter et al., 2022). The sex/gender alignment scores from the Youth Self-Report Gender Questionnaire rely on self-reported responses. Males, even from preadolescence, might feel pressured to answer gender-related questions less truthfully. Consequently, males with rsFC brain patterns less similar to the typical male pattern might still report high sex/gender alignment scores. Thus, the groups of males with higher sex/gender alignment scores may include participants whose sex and gender are not actually aligned. Two observations reinforce this possibility: 1) There were significantly more aligned males than females in the sample (1,172 vs. 873; Table 1), and 2) unaligned females had overall lower Youth Self-Report Gender Questionnaire scores compared to unaligned males (Supplementary Figure 1), both patterns also observed in the entire ABCD dataset (Potter et al., 2022). A question is thus raised of whether the Youth Self-Report Gender Questionnaire is measuring gender equally effectively for both males and females.

While rsFC successfully captured the relationship between brain patterns and sex/gender alignment in both males and females, cortical thickness only showed this relationship in males, reflecting the same negative correlation. The lack of a significant correlation between the cortical thickness sex SVM and sex/gender alignment scores in females underscores the cortical thickness’s inability to effectively capture the relationship between adolescents’ brain profiles and sex/gender alignment.

### 4.5. Functional connectivity and neuroanatomy do not predict sex/gender alignment

Neither the rsFC, cortical thickness, nor combined rsFC/cortical thickness SVR models were able to capture the variance in sex/gender alignment. A recent study by Dhamala et al. (2024) demonstrated that sex/gender alignment, as measured by the Parent-Report Gender Questionnaire, could be predicted using rsFC and linear ridge regression. Although both our study and theirs used the ABCD dataset, our findings do not replicate theirs. To assess the robustness of our findings, we also implemented linear ridge regression using online scripts provided by Dhamala et al. (2024). Similar to their approach, separate models were constructed for males and females. rsFC was used as the predictor and sex/gender alignment scores were used as labels. Our robustness check results further confirmed that rsFC is unable to predict sex/gender alignment, as measured by the Youth Self-Report and Parent-Report Gender Questionnaires. The divergence in findings could be attributed to several differences in our analyses including the use of different cortical (400 (Schaefer et al., 2018) vs. 333 (Gordon et al., 2016)) and noncortical (19 (Fischl et al., 2002) vs. 61 (Seitzman et al., 2020)) parcels, and choices that led to different final ABCD samples (e.g., Dhamala et al. (2024) excluded individuals with less than 4 minutes of data – final sample *n* = 4,757, while we excluded subjects with less than 8 minutes of data – final sample *n* = 3,196). Additionally, they band-pass filtered at 0.009 Hz ≤ f ≤ 0.08 Hz while we did at 0.008 Hz < f < 0.10 Hz.

Although rsFC effectively captures the relationship between sex and gender (significant accuracy difference when predicting aligned vs. unaligned individuals), the inability to straightforwardly predict sex/gender alignment from brain biomarkers suggests that gender may be a more complex construct that is not as clearly reflected in functional connectivity or cortical thickness patterns. Another possibility would be that linear regression models may not be able to capture potential non-linear gender effects. Additionally, the challenge of capturing variance in gender could be attributed to the overall scarcity of population-level variability in gender scores. It is possible that developmental factors play a role in this phenomenon, and the results might differ if we were examining a population at a more advanced pubertal stage. The inability to replicate findings from a study that reported sex/gender alignment prediction from rsFC using the same dataset suggests that more research is necessary to confirm the robustness and reliability of neural predictors of gender versus sex.

### 4.6. Limitations

This study has certain limitations that should be acknowledged. First, sex was assessed using a binary parent-report question. It is important to acknowledge that existing evidence and theory suggest that binary sex classification may be suboptimal, since human brains are largely composed of unique mosaics of female-typical and male-typical features (Fine, 2014; Joel, 2020). Second, the self-reported gender data displayed minimal variability. This may have contributed to our inability to detect the relationship between functional connectivity, neuroanatomy, and the degree of sex/gender alignment. Future work should include individuals with varying levels of gender nonconformity to enhance the ability to capture these relationships. Third, there is a possibility that the Youth Self-Report Gender Questionnaire may not be measuring gender equally effectively for both males and females. This is based on the different score distribution between males and females which could be due to greater social tolerance to nonconformity in young females compared to males (Potter et al., 2021; Potter et al., 2022). Future studies need to consider a measuring tool that captures gender identity, dysphoria, and expression more effectively for both sexes. Fourth, gender is shaped significantly by social and cultural factors. Since the ABCD dataset was collected entirely in the United States and is not representative of the world population (Ricard et al., 2022) future research should explore whether the relationship between brain and gender is consistent across different cultures. Lastly, this study did not provide a detailed evaluation of subcortical and cerebellar networks. Subsequent analyses should focus on assessing functional and neuroanatomical sex differences in these networks, as they are as crucial as the cortex for understanding human behavior and function.

## 5. Conclusions

In summary, our study demonstrates that brain functional connectivity (i.e., rsFC) captures the relationship between sex and gender more effectively compared to cortical thickness, as well as the extent to which sex/gender alignment correlates with a brain sex profile. While we observed normative sex differences in both functional connectivity and cortical thickness in youth, functional connectivity proved to be a more sensitive predictor of sex, with association networks playing a central role in these differences. Additionally, we found limitations in using functional connectivity and neuroanatomy to predict sex/gender alignment, highlighting the complexity of these concepts. Our findings challenge binary models of brain-sex differences, advocating for more inclusive and nuanced approaches to studying the relationship between gender and neurobiology.

## CRediT authorship contribution statement

**Athanasia Metoki:** Conceptualization, Methodology, Validation, Formal analysis, Investigation, Data Curation, Writing – Original Draft, Writing – Review & Editing, Visualization, Supervision **Roselyne Chauvin:** Writing – Review & Editing, **Evan M. Gordon:** Writing – Review & Editing, **Timothy O. Laumann:** Writing – Review & Editing, **Benjamin P. Kay:** Writing – Review & Editing, **Samuel R. Krimmel:** Writing – Review & Editing, **Scott Marek:** Writing – Review & Editing, **Anxu Wang:** Writing – Review & Editing, **Andrew N. Van:** Writing – Review & Editing, **Noah J. Baden:** Writing – Review & Editing, **Vahdeta Suljic:** Writing – Review & Editing, **Kristen M. Scheidter:** Writing – Review & Editing, **Julia Monk:** Writing – Review & Editing, **Forrest I. Whiting:** Writing – Review & Editing, **Nadeshka J. Ramirez-Perez:** Writing – Review & Editing, **Deanna M. Barch:** Methodology, Writing – Review & Editing, Supervision, **Aristeidis Sotiras:** Methodology, Writing – Review & Editing, Supervision, **Nico U.F. Dosenbach:** Methodology, Writing – Review & Editing, Supervision.

## Acknowledgments

This work was supported by NIH grants T32 MH100019 (A.M.), MH122066 (E.M.G.), MH121276 (E.M.G.), NS129521 (E.M.G.), MH129616 (T.O.L.), NS123345 (B.P.K.), MH121518 (S.M.), U01DA041120-01 (D.M.B.), MH096773 (N.U.F.D.), MH122066 (N.U.F.D.), MH121276 (N.U.F.D.), MH124567 (N.U.F.D.), NS129521 (N.U.F.D.), and NS088590 (N.U.F.D.); by the National Spasmodic Dysphonia Association (E.M.G.); by the Taylor Family Institute for Innovative Psychiatric Research (T.O.L.); by the Intellectual and Developmental Disabilities Research Center (N.U.F.D.); by the Kiwanis Foundation (N.U.F.D.); by the Washington University Hope Center for Neurological Disorders (E.M.G. and N.U.F.D.); and by Mallinckrodt Institute of Radiology pilot funding (N.U.F.D.).

