## SupplementaryMaterials for "Brain functional connectivity, but not neuroanatomy, captures the interrelationship between sex and gender in preadolescents"

**Supplementary Materials**

### 1. Supplementary Methods

#### *1.1. Support Vector Machine Classification & Regression*

The MATLAB (The MathWorks Inc., 2023) fitclinear function was used to construct the univariate and multivariate models that classified participants as male or female. Here, the notation ("option name," "option choice") is used to describe the choices made when using the fitclinear function. Support vector machine (SVM) ("Learner," "svm") was chosen for the binary classification model. Regularization was performed with the ridge (L2-norm) penalty ("Regularization," "ridge"). Dual stochastic gradient descent for SVM ("Solver," "dual") was the objective function minimization technique. Lastly, grid search optimization of the regularization coefficient C was performed. This optimization attempts to minimize the cross-validation loss (error) determining the balance between the training errors and the generalizability of the SVM classification model. Specifically, it searches for the optimum among a given C range. In this case, the range of C was [$2^{-10}$, $2^{-8}$, …, $2^{12}$, $2^{14}$] (i.e., 13 values in total) (Hsu et al., 2003). Inner accuracies were calculated for each C value and the C with the highest mean inner prediction accuracy was chosen as the optimal C (Cui & Gong, 2018; Hsu et al., 2003) for testing in each of the outer fold test sets. The support vector machine regression (SVR) models were constructed using MATLAB's fitrlinear function, with the same parameters and choices as those applied in the SVM classification models.

#### 2. Supplementary Results

##### *2.1. Cortical thickness SVM classification at the vertex-level*

We performed SVM classification to examine the predictability of sex in adolescents. We used cortical thickness at the vertex-level (59,412 cortical vertices for each participant) in an effort to capture regional effects that may not align with ROI boundaries. The cortical thickness vertex-level sex classifier was trained on the same aligned participants as the rsFC and ROI-level cortical thickness classifiers. It correctly separated aligned males from females with an accuracy of 79% ($p < 0.001$). Sensitivity, specificity, AUC, and MCC of the model were 0.77, 0.83, 0.87, and 0.59 respectively. Variation in the anatomical organization of cortical vertices belonging to the medial visual, visual, premotor, context association, and frontoparietal networks, in that order, contributed the most to the model and were therefore relatively more important in predicting participant sex. The vertex-level cortical thickness sex classifier performed significantly worse in predicting aligned individuals compared to the rsFC classifier (McNemar's test: $\chi^2 = 4.03$, $p = 0.04$). However, its performance did not significantly differ from that of the ROI-level cortical thickness classifier ($\chi^2 = 1.55$, $p = 0.21$). These findings highlight the superior performance of the rsFC classifier in predicting sex while emphasizing the limitations of the vertex-level cortical thickness classifier. Additionally, they point to the broader limitations of cortical thickness as a predictor of sex.

Next, we assessed the classifier's efficacy in predicting the sex of individuals with sex/gender unalignment. The vertex-level cortical thickness sex classifier trained on the aligned participants was able to classify unseen participants with sex/gender unalignment as male or female with 76% accuracy ($p < 0.001$; Sensitivity, specificity, AUC, MCC = 0.84, 0.72, 0.87, and 0.53). However, the vertex-level cortical thickness SVM model did not achieve a statistically significantly higher prediction accuracy for the aligned independent testing set (79%) compared to the unaligned group (76%; $z = 1.35$, $p = 0.18$).

The vertex-level cortical thickness sex classifier did not show significantly better performance in predicting unaligned individuals compared to the rsFC classifier (McNemar's test: $\chi^2 = 1.30$, $p = 0.25$). It did however show significantly better performance compared to the ROI-level cortical thickness classifier ($\chi^2 = 6.10$, $p = 0.01$).

Lastly, we investigated the relationship between the vertex-wise cortical thickness scores (brain profile predictions) and sex/gender youth-self reported and parent-reported sex/gender alignment scores to evaluate the consistency between the adolescents' brain profiles and sex/gender alignment. The correlations between vertex-level cortical thickness classification and sex/gender alignment scores were not significant for females, but were significant for males ($\rho = -0.17$, $p < 0.001$) exhibiting the same inverse relationship as in the rsFC and ROI-level cortical thickness patterns.

*2.2. Cortical thickness SVR at the vertex-level*

Following the SVM classification, we assessed whether an adolescent's degree of sex/gender alignment can be predicted by vertex-level cortical thickness. The SVM classification was extended to examine linear patterns among features that predict a continuous variable (here, sex/gender alignment as assessed by the Youth Self-Report and Parent-Report Gender Questionnaires) with linear SVR. Using the same parameters/choices as in the SVM classification models, we ran a total of four models, utilizing different combinations of features and questionnaire types: One model exclusive to females and one exclusive to males, each utilizing the Youth Self-Report Gender Questionnaire as the target variable in one iteration and the Parent-Report Gender Questionnaire in another. Similar to the rsFC and ROI-level cortical thickness SVR models, neither vertex-level sex/gender alignment SVR model successfully predicted the sex/gender alignment Youth Self-Report and Parent-Report Gender Questionnaire scores. Correlations between the original sex/gender alignment scores and predicted sex/gender alignment scores were not significant (all *p*-values > .05). and all coefficients of determination were negative ($R^2 < 0$) suggesting that the models performed worse than a null model predicting the mean of the dependent variable (sex/gender alignment scores) for all observations.

##### 3. Supplementary Figures

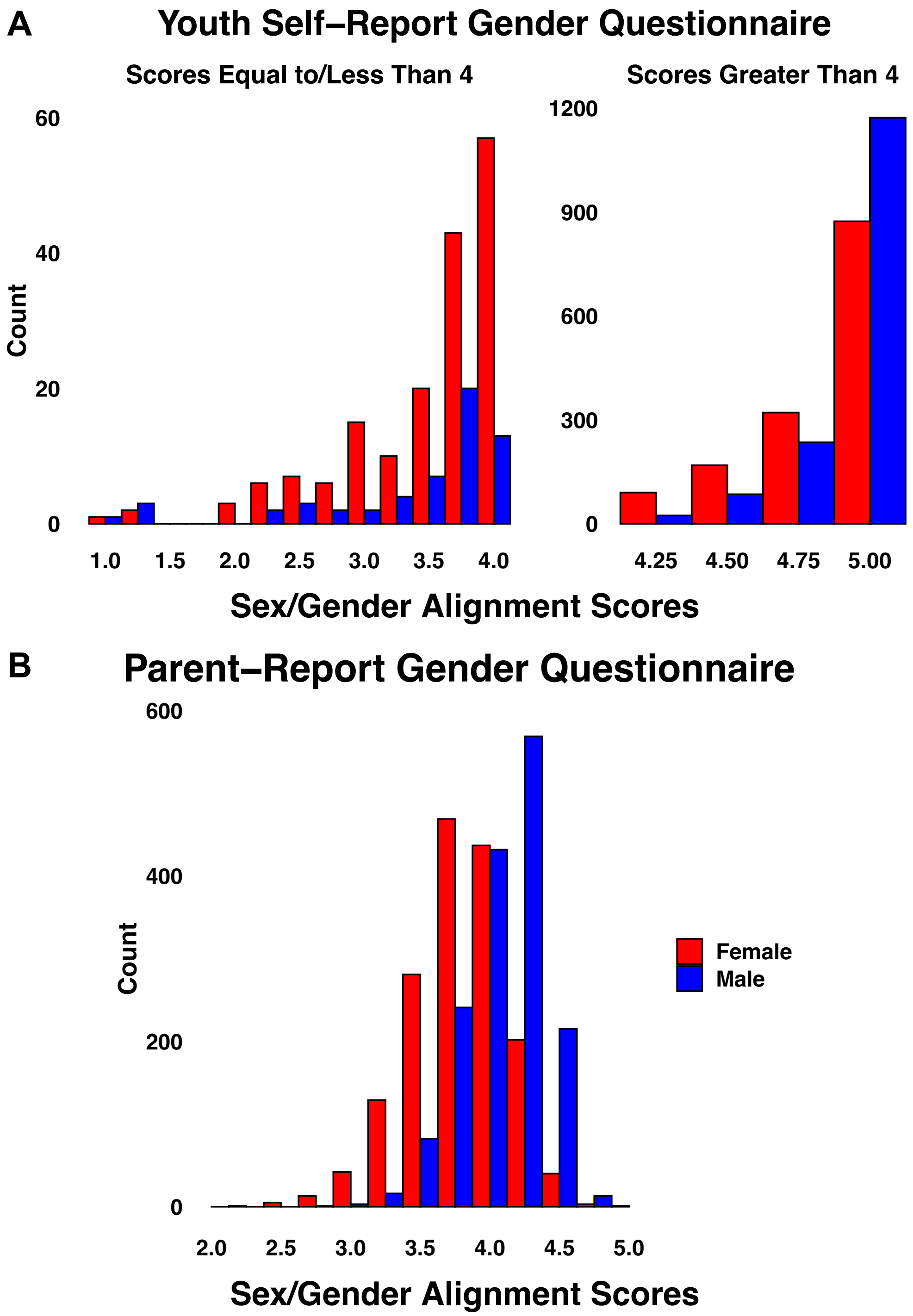


**Supplementary Figure 1.** Frequency distribution of the (A) Youth Self-Report and (B) Parent-Report Gender Questionnaires.

#### 4. Supplementary Tables

**Supplementary Table 1. ABCD Gender Survey Questions.**

| Survey | Item | Construct |
|---|---|---|
| Youth Self-Report Gender Identity Questionnaire | How much do you feel like a <boy/girl>? | Sex-congruent felt-gender |
| | How much do you feel like a <girl/boy>? | Sex-incongruent felt-gender |
| | How much have you had the wish to be a <girl/boy>? | Gender non-contentedness |
| | How much have you dressed or acted as a <girl/boy> during play? | Gender non-conformity |
| Parent-Report Gender Identity Questionnaire | <His/Her> favorite playmates are: | Sex-typed behavior during play |
| | <He/She> plays with girl-type dolls, such as “Barbie”. | |
| | <He/She> plays with boy-type dolls such as action figures or “GI-Joe”. | |
| | <He/She> experiments with cosmetics (makeup) and jewelry. | |
| | <He/She> imitates female characters seen on TV or in the movies. | |
| | <He/She> imitates male characters seen on TV or in the movies. | |
| | <He/She> plays sports with boys (but not girls). | |
| | <He/She> plays sports with girls (but not boys). | |
| | *In playing “mother/father”, “house”, or “school” games, <he/she> takes the role of:* | |
| | <He/She> plays “girl/boy-type” games (as compared to “boy/girl-type” games). | |
| | *In dress-up games, <he/she> likes to dress up as:* | |
| | <He/She> states the wish to be a <girl/boy> or <woman/man>. | Gender dysphoria |
| | <He/She> states that <he/she> is a <girl/boy> or <woman/man>. | |
| | <He/She> talks about not liking <his/her> sexual anatomy (private parts). | |

**Supplementary Table 2. Family income and descriptive statistics.**

| | TOTAL (n = 3,196) | | |
|---|---|---|---|
| | Males<br>n = 1,573; 49.2% | Females<br>n = 1,623; 50.8% | |
| Family Income | | | |
| < $5,000 | 32 (2%) | 26 (1.6%) | $\chi^2 = 0.84$, p = 0.36 |
| $5,000 – $11,999 | 28 (1.8%) | 34 (2.1%) | $\chi^2 = 0.42$, p = 0.52 |
| $12,000 – $15,999 | 27 (1.7%) | 23 (1.4%) | $\chi^2 = 0.46$, p = 0.50 |
| $16,000 – $24,999 | 51 (3.2%) | 51 (3.1%) | $\chi^2 = 0.03$, p = 0.87 |
| $25,000 – $34,999 | 85 (5.4%) | 78 (4.8%) | $\chi^2 = 0.59$, p = 0.44 |
| $35,000 – $49,999 | 97 (6.2%) | 108 (6.7%) | $\chi^2 = 0.32$, p = 0.57 |
| $50,000 – $74,999 | 183 (11.6%) | 196 (12.1%) | $\chi^2 = 0.15$, p = 0.70 |
| $75,000 – $99,999 | 215 (13.7%) | 225 (13.9%) | $\chi^2 = 0.03$, p = 0.87 |
| $100,000 – $200,000 | 534 (33.9%) | 572 (35.2%) | $\chi^2 = 0.59$, p = 0.44 |
| > $200,000 | 223 (14.2%) | 220 (13.6%) | $\chi^2 = 0.26$, p = 0.61 |

#### 5. Supplementary Equations

$$MCC = \frac{Cov(c,l)}{\sigma_c * \sigma_l} = \frac{TP * TN - FP * FN}{\sqrt{(TP + PF) * (TP + FN) * (TN + FP) * (TN + FN)}}$$

**Supplementary Equation 1. Matthews Correlation Coefficient.** Measures the correlation of the true classes *c* with the predicted labels *l*. Worst value = -1; Best value = +1. *Cov*(*c*,*l*): covariance of the true classes *c* and predicted labels *l*; $\sigma_c$: standard deviation of the true classes; $\sigma_l$: standard deviation of the predicted labels; TP: True Positives; TN: True Negatives; FP: False Positives; FN: False Negatives.
